# MOSAIC enables *in situ* saturation mutagenesis of genes and CRISPR prime editing guide RNA optimization in human cells

**DOI:** 10.1101/2024.04.25.591078

**Authors:** Jonathan Y. Hsu, Kin Chung Lam, Justine Shih, Luca Pinello, J. Keith Joung

## Abstract

CRISPR prime editing offers unprecedented versatility and precision for the installation of genetic edits *in situ*. Here we describe the development and characterization of the Multiplexing Of Site-specific Alterations for *In situ* Characterization (**MOSAIC**) method, which leverages a non-viral PCR-based prime editing method to enable rapid installation of thousands of defined edits in pooled fashion. We show that MOSAIC can be applied to perform *in situ* saturation mutagenesis screens of: (1) the *BCR-ABL1* fusion gene, successfully identifying known and potentially new imatinib drug-resistance variants; and (2) the *IRF1* untranslated region (UTR), re-confirming non-coding regulatory elements involved in transcriptional initiation. Furthermore, we deployed MOSAIC to enable high-throughput, pooled screening of hundreds of systematically designed prime editing guide RNA (**pegRNA**) constructs for a large series of different genomic loci. This rapid screening of >18,000 pegRNA designs identified optimized pegRNAs for 89 different genomic target modifications and revealed the lack of simple predictive rules for pegRNA design, reinforcing the need for experimental optimization now greatly simplified and enabled by MOSAIC. We envision that MOSAIC will accelerate and facilitate the application of CRISPR prime editing for a wide range of high-throughput screens in human and other cell systems.

## Introduction

Prime editing (**PE**) is a recently described gene-editing technology that can install essentially any DNA base substitution or short insertion/deletion mutation^1^, thereby greatly expanding the repertoire of genetic alterations that can be introduced at target genomic loci relative to CRISPR-Cas nucleases^2,3,4^ and base editors (**BE**)^5,6,7^. Prime editors have two components: (1) a prime editor 2 (**PE2**) protein, which is a fusion of a *Streptococcus pyogenes* Cas9 (SpCas9) nickase and an engineered Moloney murine leukemia virus reverse transcriptase (**MMLV-RT**) domain, and (2) a prime editing guide RNA (**pegRNA**), which consists of a standard SpCas9 guide RNA and additional 3’ elements in the primer binding site (**PBS**) and reverse transcription template (**RTT**) sequences, the latter bearing the edit of interest (**Extended Data Fig. 1a**). The prime editor-pegRNA complex introduces a nick into the non-target strand at the target site and the resulting liberated DNA strand anneals to the PBS element of the pegRNA. The MMLV-RT then reverse transcribes the adjacent RTT element to create a DNA flap encoding the desired edit (**Extended Data Figs. 1b-1c**). Incorporation of this newly synthesized, edit-encoding DNA flap into the genomic locus results in the intended genetic alteration (**Extended Data Figs. 1d – 1e**). Addition of a nicking sgRNA (**ngRNA**), referred to as the PE3 strategy^1^, introduces a nick on the opposing target DNA strand and can bias repair of the unedited target strand to thereby increase editing frequency (**Extended Data Figs. 1d-1e**).

Despite its unprecedented capability to make essentially any small genetic edit of interest, certain characteristics of the prime editing system make it more challenging to use for potential research and therapeutic applications. Optimization of pegRNAs for introducing any given individual desired edit can be complex because the lengths of the PBS and RTT sequences have been shown to influence prime editing efficiencies^1^. Therefore, optimization of pegRNAs would ideally include pairwise combinatorial testing of a large number of various PBS and RTT lengths. An additional challenge for library screening is that prime editing efficiencies (especially in cell types other than HEK293s) are generally much lower than those achieved with CRISPR nuclease or base editor technologies, even with recent advances that include pegRNA modifications and co-expression of a dominant-negative protein involved in DNA mismatch repair^8,9^.

Here we describe development, validation, and applications of the Multiplexing Of Site-specific Alterations for *In situ* Characterization (**MOSAIC**) method. This approach is based on high-throughput screening with pooled non-viral pegRNA libraries (“**pegPools**”) encoded on linear DNA fragment cassettes that can be rapidly constructed by PCR, with sequence diversity introduced using mixed base or high-throughput oligonucleotide synthesis (**Fig. 1a**, top panel; **Supplementary Note 1**). We show that MOSAIC can be used successfully to perform saturation mutagenesis of coding and non-coding endogenous sequences in human cells and to optimize the efficiencies of prime editors by comprehensively testing pairwise combinations of various length RTT and PBS sequences.

**Figure 1:**
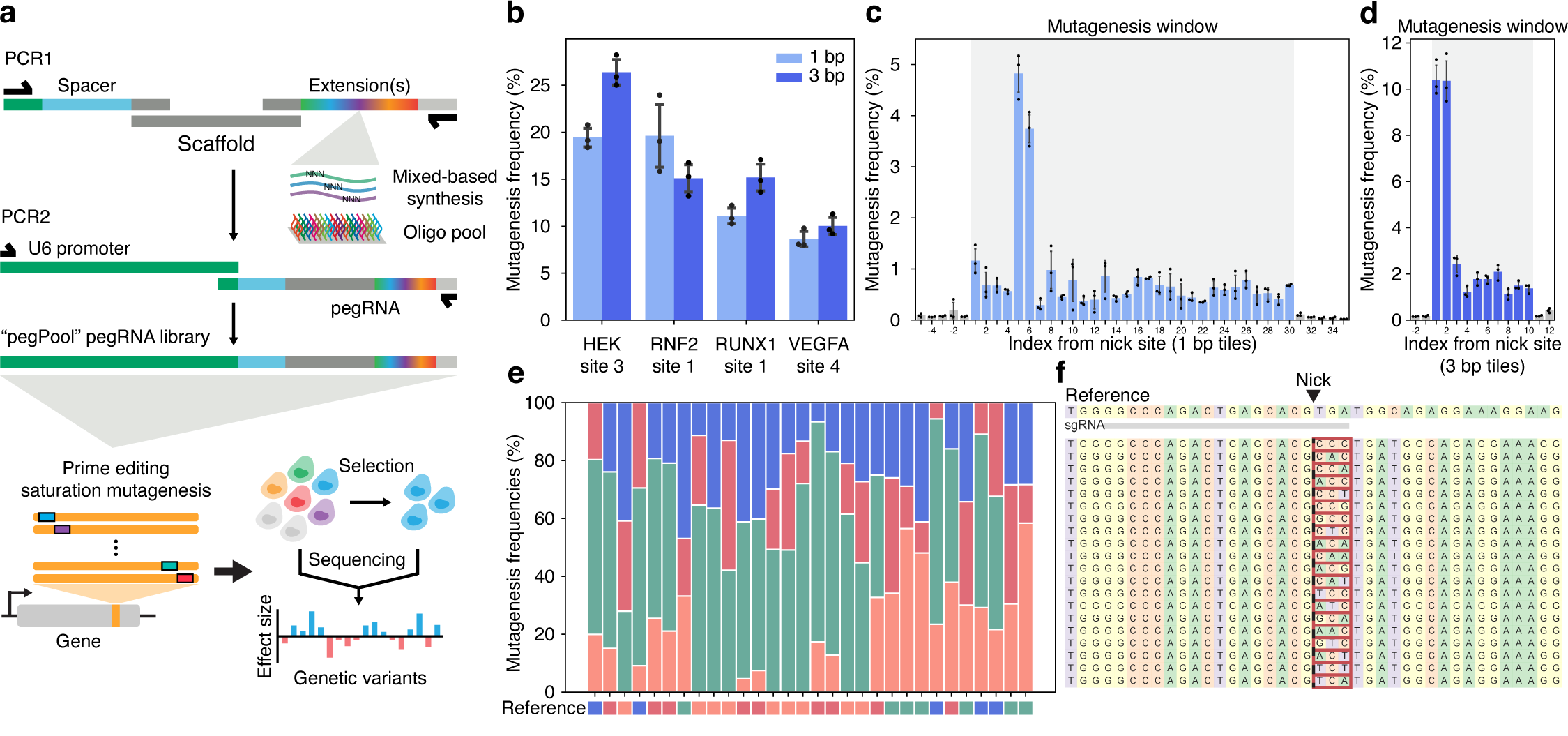
MOSAIC method for pooled prime editing in human cells. **(a)** Schematic of PCR protocol for the rapid generation of pooled pegRNA constructs (pegPools) (top) and *in situ* saturation mutagenesis with prime editing (bottom). **(b)** Pooled prime editing efficiencies for the installation of single– or triple-base saturation mutagenesis across four endogenous gene targets in HEK293T cells. The data are meanlJ±lJs.d. of nlJ=lJ3 independent biological replicates. **(c,d)** Saturation mutagenesis across HEK site 3 for the installation of single-base and triple-base substitutions. The data are meanlJ±lJs.d. of nlJ=lJ3 independent biological replicates. **(e)** Proportion of substitution edits for single-base saturation mutagenesis at HEK site 3. **(f)** Visualization of the top allelic outcomes with a MOSAIC pooled library encoding randomized insertion edits of length 3.

## Results

An initial application we envisioned for MOSAIC was to perform saturation mutagenesis of various endogenous genomic loci. Because PE can introduce all possible base changes at any given genomic locus, an appropriately designed pegPool library could in principle induce full randomization of either codons or individual DNA bases within gene coding sequences or non-coding regulatory sequences, respectively (**Fig. 1a**, bottom panel). Such a pegPool library might then be introduced together with other components needed for PE into human cells in culture and then the resulting population of modified cells could be assessed for desired phenotypes of interest either by screening or selection. Cells with the phenotype (and potentially those without) could subsequently be sequenced to identify associated genomic edits at the randomized locus (**Fig. 1a**, bottom panel).

To assess the robustness of MOSAIC for performing *in situ* saturation mutagenesis of endogenous genes in cells, we built pegPool libraries designed to install comprehensive single-base or codon triplet substitution edits across 30 bp sequence windows at four different human genomic loci (**Methods; Supplementary Table 1**). Following introduction of each of these eight different pegPool libraries with plasmids expressing PE2 protein and a corresponding ngRNA into HEK293T cells, we found mean total editing frequencies (i.e. sum of all potential edits encoded by each library) ranging from 8.63% to 26.38% for each library (**Fig. 1b**). These edit frequencies are comparable to what has been previously reported with transfection of plasmids encoding pegRNAs into these same cells, suggesting that transfection of pegPool DNA fragments provides an efficacious strategy for expressing pegRNAs. Analysis of sequencing data at each target site showed that the alterations introduced were relatively uniform across the target editing windows, with the exception of positions within the protospacer adjacent motif (**PAM**) where we observed significantly higher editing frequencies (**Figs. 1c-1d**; **Extended Data Figs. 2a-2f**). One potential reason for this finding is that re-targeting and re-editing (with reversion of earlier mutations introduced earlier) may occur at the target site unless or until the canonical bases of the PAM become mutated. In addition, at each position within the editing windows, we observed representation of all three potential alternative bases introduced by each pegPool library but with some preference for replacement with a cytosine (**Fig. 1e**). We further tested whether pegPools could be used to introduce short, randomized insertions of lengths 3, 6, and 9 bp and found that with these libraries we were able to install as many as 10,000 unique insertion edits into the genome in a single transfection experiment (**Fig. 1f**, **Extended Data Figs. 3a-3b**, **Supplementary Table 1**).

Having established the efficacy of using transfected pegPool libraries to express pegRNAs in human cells, we performed a proof-of-principle test by using MOSAIC to perform saturation mutagenesis of a region of the *BCR-ABL1* gene translocation coding sequence (**Fig. 2**), an oncogenic driver of chronic myeloid leukemia (**CML**) cells^10^. Previous studies have shown that specific mutations within the *ABL1* tyrosine kinase gene part of this translocation can confer drug-resistance to imatinib, a tyrosine kinase inhibitor used as a first-line therapy in *BCR-ABL1*-positive CML patients^11^. Our screening of 84 different pegRNA and ngRNA combinations targeting this imatinib binding site region yielded several pegRNA-ngRNA combinations that could mediate efficient mutagenesis of this sequence in HEK293T cells (**Extended Data Fig. 4**). Using a pegRNA that could efficiently mutagenize the imatinib-binding site (amino acids 311-318) within the *BCR-ABL* sequence (**Extended Data Fig. 4**, **Supplementary Table 2**), we constructed a pegPool library designed to induce codon saturation mutagenesis of this region (**Methods**). We electroporated this pegPool library together with plasmids encoding PE2 and an accompanying efficiency-enhancing ngRNA (**Extended Data Fig. 5**, **Supplementary Table 2**) into K562 cells, a CML patient-derived cell line that harbors an endogenous *BCR-ABL1* gene and that is sensitive to imatinib^12^. After 3 days, transfected K562 cells were split into two cultures in duplicate – one set treated with DMSO (negative control) and the other with imatinib (**Fig. 2a**). After 7 additional days, we performed targeted amplicon sequencing of *BCR-ABL1* complementary DNA (cDNA) prepared from the mRNA of edited cells from each culture condition (**Fig. 2a**). For both conditions, we observed complete coverage of NNK codons encoding every amino acid variant across all targeted *BCR-ABL1* codons (**Extended Data Figs. 5a-5g**; **Extended Data Figs. 6a-6d**). However, genetic variation at residue 315 was significantly enriched in the imatinib-treated samples compared to the DMSO-treated controls (P<0.01) (**Fig. 2b**) for specific amino acids, including the T315I “gatekeeper” mutation commonly identified in patients with imatinib-resistant CML (p<0.01) (**Fig. 2c**; **Extended Data Fig. 7**; **Supplementary Table 3**). Interestingly, we also observed several other mutations at the same amino acid position (T315M, T315L, and T315E) that were significantly enriched in the imatinib-treated samples (P<0.01) (**Fig. 2c**). Although two of these mutations have been previously observed in an *in vitro* cell culture study designed to identify imatinib resistance mutations^13^, none have been reported from imatinib-resistant CML patient samples. One potential reason for this is that obtaining these amino acid substitutions requires at least two base mutations to the wild-type codon (in contrast to the single base mutation required for the T315I substitution) and therefore would be less likely to occur in patients.

**Figure 2:**
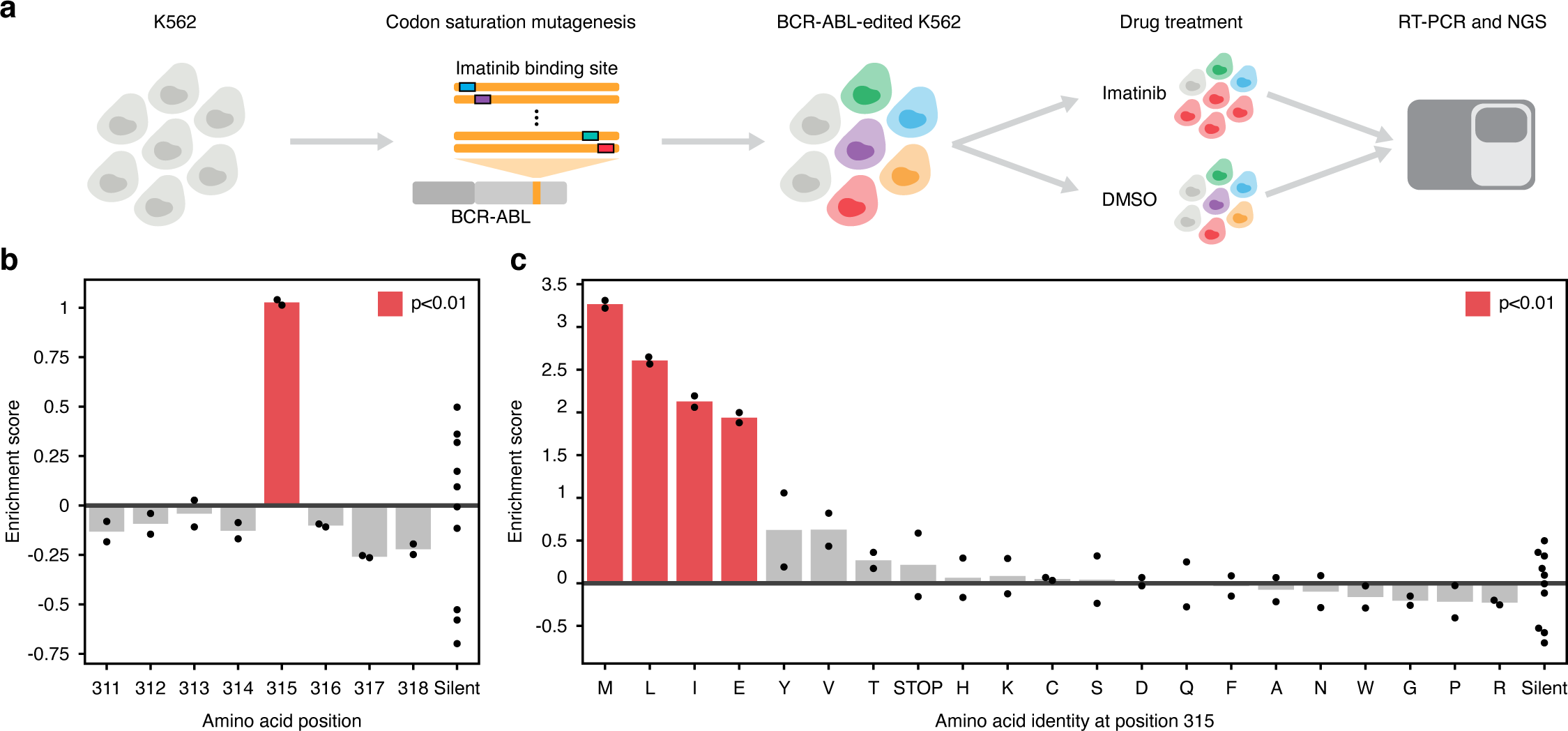
Application of MOSAIC for *in situ* saturation mutagenesis of the *BCR-ABL1* translocation gene in human cells. **(a)** Schematic for *in situ* functional screening of genetic variants associated with imatinib resistance in K562 cells. **(b)** Enrichment scores of amino acid substitutions within the BCR-ABL1 imatinib binding site associated with imatinib resistance. Experiments shown were performed in two independent biological replicates. **(c)** Enrichment scores of amino acid variants at position 315 associated with imatinib resistance. Experiments shown were performed in two independent biological replicates.

In addition to altering coding sequences, we sought to test if MOSAIC could be used to perform saturation mutagenesis of non-coding elements. To do this, we used MOSAIC to comprehensively mutate non-coding regulatory sequences in the 5’ untranslated region (**UTR**) of the transcription factor gene *IRF1* (**Extended Data Fig. 8a**). Previous studies have defined a core downstream promoter element (**DPE**) that positively influences transcription initiation from the *IRF-1* promoter^14^. Using a pegRNA we identified as being efficient for editing a 36 bp window that overlaps this DPE, we constructed a pegPool library that fully randomizes bases across that window in three base pair sub-windows (**Extended Data Fig. 8b**; **Supplementary Table 4**). Following transfection of this pegPool library and plasmids encoding an accompanying ngRNA and PE2 fusion protein into HEK293T cells, we performed targeted amplicon sequencing of both the genomic DNA and cDNA reverse transcribed from RNA isolated from the edited cells (**Extended Data Figs. 8a-8b**). Comparing the proportions of variants we found on cDNA relative to those found on genomic DNA, we observed statistically significant depletion of two contiguous 3 bp tiles across the entire mutagenesis window (p<0.01) (**Extended Data Fig. 8c**). Notably, these depleted tiles directly overlap the consensus motif of the DPE (**Extended Data Fig. 8c**, orange colored text), confirming that MOSAIC can also be used to mutagenize and fine map non-coding regulatory elements *in situ* at high resolution.

We additionally used MOSAIC for the completely different application of optimizing the editing activities of pegRNAs. As noted above, the lengths of the PBS and RTT segments within a pegRNA can potentially influence prime editing activity^1,15,16^, but testing the large matrix of different PBS and RTT length combinations can be labor-intensive if tested experimentally one-by-one. To address this challenge, we envisioned that for any given desired edit, we could design a pegPool comprised of pegRNAs with various combinations of PBS and RTT length, each barcoded with a unique 4 bp insertion edit (**Fig. 3a**). We further reasoned that the installation frequencies of these 4-mer barcodes mediated by each pegRNA in human cells would provide quantitative data to identify the most efficient PBS/RTT combinations among those tested in the pool (**Fig. 3a**).

**Figure 3:**
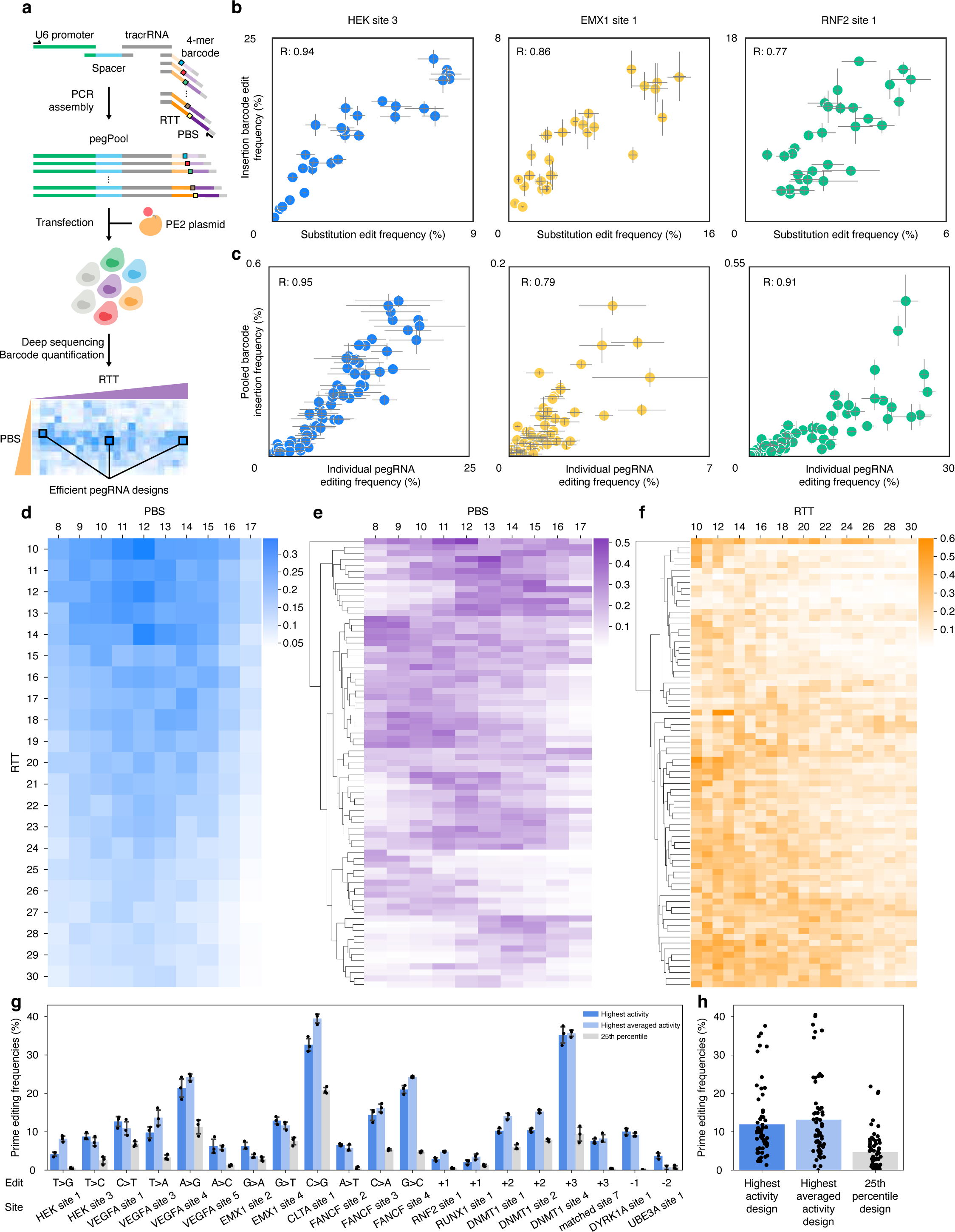
Application of MOSAIC for pegRNA activity optimization in human cells. **(a)** Schematic overview illustrating the use of MOSAIC for high-throughput pooled screening of pegRNA designs. **(b)** Scatterplots of prime editing efficiencies for installing a 4-mer insertion barcode (y-axes) or substitution edit (x-axes) with various pegRNA designs for three different endogenous gene target sites in HEK293T cells. Data shown are from three independent biological replicates, meanslJ±lJs.d. are indicated. **(c)** Scatterplots of prime editing efficiencies when installing an insertion barcode with pegRNAs in pooled format (y-axes) or individually (x-axes) targeted to three different endogenous gene target sites in HEK293T cells. Correlation (R) was determined with Spearman’s Rho. Data shown are from three independent biological replicates, meanslJ±lJs.d. are indicated. **(d)** Averaged enrichment scores for 18,690 pegRNA designs for 210 different combinations of various length PBS (x-axis) and RTT (y-axis) sequences across 89 different target sites. Heat map squares reflect relative enrichment. **(e)** Averaged PE efficiencies for 89 target sites (y-axis) for 18,690 pegRNAs across 10 different length PBS sequences (x-axis). Heat map squares reflect relative enrichment scores. **(f)** Averaged PE efficiencies for 89 target sites (y-axis) for 18,690 pegRNAs across 21 different length RTT sequences (x-axis). Heat map squares reflect relative enrichment scores. **(g)** Individual testing of pegRNA designs derived from MOSAIC pegRNA optimization experiments for 20 different target gene loci in human HEK293T cells. Data shown are from three independent biological replicates, meanslJ±lJs.d. are indicated. **(h)** Averaged prime editing efficiencies of different types of pegRNA designs tested in (**g**). Data shown are from three independent biological replicates, meanslJ±lJs.d. are indicated.

To test the feasibility of this envisioned strategy, we performed two control experiments. First, we compared the efficiencies of pegRNAs installing 4-mer barcode insertions to those that introduce point mutations and observed a strong correlation in the relative activities of pegRNAs spanning 90 different designs across three different target genomic loci (**Fig. 3b**; **Supplementary Table 5**). Second, we assessed whether the relative editing efficiencies of pegRNAs tested in our pooled format recapitulated their activities when tested individually. To do this, we compared the barcode insertion frequencies of >210 pegRNAs in three pegPools targeted to different genomic sites with the editing frequencies of individual pegRNAs in HEK293T cells (**Supplementary Table 6**) and found a strong correlation between the pooled and individual pegRNA PE efficiencies at all three sites tested (Spearman R values of 0.95, 0.79, and 0.91) (**Fig. 3c**). We also observed strong correlation between editing efficiencies of pegRNAs from our pooled pegRNA experiments and individual pegRNAs tested in a previous report^1^ (Spearman R = 0.90) (**Extended Data Fig. 9**).

Having established the efficacy of using MOSAIC to assess the relative editing efficiencies mediated by various pegRNA designs in pooled format, we next used it at larger scale to characterize libraries of different pegRNAs for 89 different target spacer sites. We constructed 89 pegPool libraries, each harboring 210 barcoded pegRNA candidates that encompassed combinations of 21 different RTT lengths (ranging from 10 to 41 nts depending on the target site) and 10 different PBS lengths (8 to 17 nts) (**Supplementary Tables 7, 8, and 9**: **Supplementary Note 2**). Following transfections of each of these 89 pegPool libraries with a PE2 expression plasmid into HEK293T cells, we performed targeted amplicon sequencing to assess the relative installation frequencies of the different barcodes from each pool at the target site (**Fig. 3a**). A high-level view of the averaged relative activities of 15,960 pegRNAs from 76 target spacers (210 pegRNAs per target spacer all with RTT lengths ranging from 10 to 30 nts and PBS lengths ranging from 8 to 17 nts) suggests that those with combinations of 12/10, 12/12, and 12/14 nts, respectively, have the highest activities (**Fig. 3d, Extended Data Fig. 10**). However, when looking at activities of the pegRNAs for each of these 76 target sites averaged across the 10 different PBS lengths (**Fig. 3e**) or the 21 different RTT lengths (**Fig. 3f**), it becomes apparent that no single PBS or RTT length is consistently optimal across different spacers. The full diversity of optimal PBS/RTT lengths for different target sites can be seen by examining the activities of the 210 different pegRNA designs for each of the 89 spacer targets (**Extended Data Fig. 11**).

To further confirm the efficacy of MOSAIC for identifying optimized pegRNA designs, we used data from our pooled screens to design and individually test pegRNAs in human cells. For each of 20 different target sites, we constructed three different pegRNAs harboring: (1) the highest activity PBS/RTT combination from the pegPool screen (from **Extended Data Fig. 11**), (2) PBS and RTT lengths showing the highest averaged activity for the target spacer (from **Figs. 3e and 3f**), and (3) the PBS/RTT combination from the pegPool screen that had activity at the 25^th^ percentile (from **Extended Data Fig. 11**). These pegRNAs were designed to introduce a range of different alterations (all possible base substitutions and a variety of small insertions or deletions) (**Fig. 3g**) and, in contrast to their counterparts in the pegPool libraries, none of them had barcodes (**Supplementary Tables 9 and 10**; **Supplementary Note 2**). Testing of these various pegRNAs with PE2 in human HEK293T cells revealed that, for each of the 20 target sites, the pegRNAs harboring the highest activity and highest averaged activity PBS/RTT length combinations performed comparably to each other and consistently outperformed the pegRNA harboring the 25^th^ percentile activity PBS/RTT length combination (**Fig. 3g**). The frequencies of prime editing observed for the most optimal pegRNAs across the 20 different target sites ranged from 3.50% to 39.5% (**Fig. 3g**), values that are generally comparable to or higher than those previously reported when using the PE2 system. On average, the pegRNAs with the highest activity and highest averaged activity PBS/RTT lengths induced editing frequencies that were approximately three-fold higher than those with the 25^th^ percentile activity PBS/RTT lengths (**Fig. 3h**). Taken together, our findings demonstrate the efficacy of MOSAIC for identifying pegRNA designs with optimized editing activities in human cells.

## Discussion

We envision that the development of MOSAIC should advance and extend the applications for CRISPR prime editing in human and other mammalian cells. These applications have already begun to be explored – for example, while this manuscript was in preparation, another report described the use of PE-based saturation mutagenesis with two disease-associated genes, *NPC1* and *BRCA2* (using plasmid-encoded pegRNAs)^17^. However, by contrast, because MOSAIC sidesteps the need for cloning into plasmid or viral vectors, pegPool libraries can be more rapidly constructed by PCR and then used to install thousands of random or defined edits. Our proof-of-principle studies in which we successfully re-identified cancer drug-resistant oncogene mutations and *cis*-acting elements that regulate gene transcription demonstrate that MOSAIC can be used to perform saturation mutagenesis of both coding regions and non-coding regulatory regions of endogenous genes in human cells. We expect that there will be many additional applications for MOSAIC-mediated saturation mutagenesis beyond those described here. For example, we have also shown how the method can be used to fine tune the activities of ectopically inserted endogenous transcription factor binding sites that are placed into target gene promoters by comprehensively mutating each base position or by interrogating collections of binding sites for different factors (an approach we call PERSIST-On; Tak and Hu et al., manuscript in preparation). Similar strategies might also be applied to decode the functions of naturally occurring transcription factor binding sites and the impacts of genetic variation on endogenous gene expression.

One current limitation of the MOSAIC method is that the low efficiency of PE in human cells imposes a requirement that the target gene either be haploid or that the desired phenotype can be screened for even if only one gene allele is altered (e.g., gain-of-function mutations). Others have recently reported a strategy for inducing a haploid state for an endogenous locus of interest and as noted above, used this for performing pooled PE screens using plasmid-based pegRNA libraries^17^. In addition, we envision that this limitation will eventually be overcome as the efficiency of prime editors are further increased with technological advancements as has been the case for other platforms such as cytosine and adenine base editors^18^.

Our application of MOSAIC to address the challenge of pegRNA architecture optimization not only demonstrates the complexity of pegRNA design but also validates the use of our method as a facile solution to address this problem. The results from our testing of 210 designs for each of 89 different target sites provide one of the largest datasets of pegRNA activities described to date – a total of 18,690 pegRNAs. The diversity of different PBS/RTT length combinations we obtained for these 89 sites definitively shows that no simple design rules exist and establishes the need to experimentally optimize these parameters for each target of interest. MOSAIC provides a simple and rapid method to do so, enabling the screening of large numbers of pegRNA designs without the need for cloning or for viral vector technologies. pegRNAs optimized by MOSAIC can be further enhanced for their prime editing efficiencies by combining them with various ngRNAs (PE3 strategy) or with other recent advances such as epegRNAs and co-expression of a dominant-negative DNA mismatch repair protein^8,9,19^. In sum, the development and availability of MOSAIC should broadly extend and enhance both the research and therapeutic applications of PE technology.

## Methods

### PCR-generated prime editing guide RNAs

Prime editing guide RNAs (pegRNAs) and nicking sgRNAs (ngRNAs) constructs used in this study were in the form of PCR products. A detailed protocol is available in Supplementary Note 1. Briefly, two sequential PCR assembly steps were performed to construct the pegRNA or ngRNA constructs. The first PCR assembly step assembles a spacer sequence oligo, tracrRNA sequence oligo, and 3’ extension sequence oligo(s) (for pegRNAs) into a product consisting of the pegRNA or ngRNA sequence. The second PCR assembly step fuses the U6 promoter sequence with the PCR product from the first assembly step to construct pegRNA or ngRNA constructs capable of expression in human cells. Standard PCR protocol was used for all reactions and PCR reactions were cleaned up using paramagnetic beads and 75% ethanol washes to isolate desired amplification products.

### Design of BCR-ABL1 and IRF1 5’ UTR mutagenesis pegRNAs

Mutagenesis of *BCR-ABL1* and *IRF1* 5’ UTR was performed with PCR-generated prime editing guide RNAs using mixed-base synthesis. For *BCR-ABL1*, oligonucleotides encoding NNK tiles were used to construct the pegRNA library to introduce all possible amino acid variants at positions 311-318. For *IRF1* 5’ UTR, oligonucleotides encoding NNN tiles were used to construct the pegRNA library to introduce diverse mutagenesis across the editing window of interest.

### Oligonucleotide library design for pegRNA optimization

An oligonucleotide library containing 21,210 members was synthesized by Agilent (Supplementary Table 8). Each library member contained a spacer sequence, tracrRNA sequence, and pegRNA 3’ extension of varied length. For each spacer sequence, a total of 210 PBS-RTT length combinations were designed for the 3’ extension with PBS lengths spanning 8-17 nt and RTT lengths typically spanning 10-30 nt, where each of these PBS-RTT length combinations were uniquely barcoded with a 4-mer insertion edit placed at the +1 position. Unique flanking sequences were used to amplify sub-libraries from the oligonucleotide pool corresponding to a specific pool of pegRNA designs for a single spacer sequence. Following the amplification of the sub-libraries, subsequent PCR reactions were used to fuse a U6 promoter element to each of the sub-libraries for expression in human cells. Purified PCR products for these sub-libraries were then used for transfection to perform pooled pegRNA optimizations.

### Human cell culture

STR-authenticated HEK293T (CRL-3216) and K562 (CCL-243) were used in this study. HEK293T cells were grown in Dulbecco’s modified Eagle medium (DMEM) (Gibco) with 10% heat-inactivated fetal bovine serum (FBS) (Gibco) supplemented with 1% penicillin-streptomycin (Gibco) antibiotic mix. K562 cells were grown in Roswell Park Memorial Institute (RPMI) 1640 Medium (Gibco) with 10% FBS supplemented with 1% pen-strep and 1% GlutaMAX (Gibco). Cells were grown at 37 °C in 5% CO_2_ incubators and periodically passaged on reaching around 80% confluency. Cell culture media supernatant was tested for mycoplasma contamination every 4 weeks using the MycoAlert PLUS mycoplasma detection kit (Lonza) and all tests were negative throughout the experiments.

### Cell transfections

HEK293T cells were seeded at 1.25lJ×lJ10^4^ cells per well into 96-well flat bottom cell culture plates (Corning) or 3lJ×lJ10^5^ cells per well into 6-well cell culture plates (Corning). Then, 24 h post-seeding, cells were transfected with 40 ng of prime editor plasmid and 13.3 ng of pegRNA PCR product (and 4.4 ng nicking sgRNA PCR product for PE3) using 0.6 µl of TransIT-X2 (Mirus) lipofection reagent for experiments in 96-well plates, or 1500 ng prime editor plasmid and 500 ng of pegRNA PCR product (and 166.6 ng nicking sgRNA PCR product for PE3) and 15 µl TransIT-X2 for experiments in 6-well plates. K562 cells were electroporated using the SF Cell Line Nucleofector X Kit L (Lonza), according to the manufacturer’s protocol with 1lJ×lJ10^6^ cells per nucleofection and 6000 ng prime editor plasmid, 1500 ng pegRNA PCR product (and 500 ng nicking sgRNA PCR product for PE3). Then, 72 h post-transfection, cells were lysed for extraction of genomic DNA (gDNA) or RNA. For the drug resistance experiments, K562 cells were treated with 1.25 µM imatinib (Sigma Aldrich) or DMSO (Sigma Aldrich) 72 h post-transfection and cultured for an additional 7 days before RNA extraction.

### DNA extraction

HEK293T cells were washed with 1× PBS (Corning) and lysed overnight by shaking at 55lJ°C with 43.5 µl of gDNA lysis buffer (100 mM Tris-HCl at pHlJ8, 200 mM NaCl, 5 mM EDTA, 0.05% SDS) supplemented with 5.25 µl of 20lJmglJml^−1^ Proteinase K (NEB) and 1.25 µL of 1 M DTT (Sigma) per well for experiments in 96-well plates, or with 435 µl DNA lysis buffer, 52.5 µl Proteinase K and 12.5 µl 1 M DTT per well for experiments in 6-well plates. K562 cells were centrifuged for 5 min, media removed and lysed overnight by shaking at 55lJ°C with 870 µl DNA lysis buffer, 105 µl Proteinase K and 25 µl 1 M DTT per well in 1.5 mL Eppendorf tubes. Subsequently, gDNA was extracted from lysates using 1–2× paramagnetic beads, washed twice with 75% ethanol, and eluted in 50-200 µl of 0.1× EB buffer. DNA extraction was performed using a Biomek FX^P^ Laboratory Automation Workstation (Beckman Coulter) when using 96-well plates.

### RNA extraction and reverse transcription

At 72 h post-transfection, total RNA was extracted from cells using the NucleoSpin RNA Plus Kit (Clontech, 740984.250). Following RNA extraction, 50–250 ng of purified RNA was used for cDNA synthesis using a High-Capacity RNA-to-cDNA Kit (Thermo Fisher, 4387406). The resulting cDNA was then used downstream as a template for PCR amplification for targeted amplicon sequencing.

### Targeted amplicon sequencing

The gDNA and cDNA concentrations of samples were measured using the Qubit dsDNA HS Assay Kit (Thermo Fisher). The first PCR was performed to amplify the regions of interest (200–250 bp) using 50–100 ng of gDNA or cDNA. Primers for PCR1 included Illumina-compatible adapter sequences (Supplementary Table 10). A synergy HT microplate reader (BioTek) was then used at 485/528 nm with the Quantifluor dsDNA quantification system (Promega) to measure the concentration of the first PCR products. PCR products from different genomic amplicons were then pooled and cleaned with 0.7x paramagnetic beads. The second PCR was performed to attach unique barcodes to each amplicon using 50–200 ng of the pooled PCR1 products and barcodes that correspond to Illumina TruSeq CD indexes. The PCR2 products were again cleaned with 0.7x paramagnetic beads and measured with the Quantifluor system before final pooling. The final library was sequenced using an Illumina Miseq (Miseq Reagent Kit v.2; 300 cycles, 2lJ×lJ150lJbp, paired-end). The FASTQ files were downloaded from BaseSpace (Illumina).

### Data analysis

Amplicon sequencing data were analyzed with CRISPResso2 v.2.0.31. Depending on position and length of the intended edit(s), parameters for CRISPResso2 analysis were modified (-w, –wc, –qwc) to center the quantification window around the intended editing window. Downstream analysis was conducted using Python 3.7.6 with data sourced from ‘Quantification_window_nucleotide_frequency_table.txt’, ‘CRISPResso_quantification_of_editing_frequency.txt’, and ‘Alleles_frequency_table.txt’. Enrichment scores were calculated using the log_2_ fold-change of normalized sample counts over control counts.

### Statistics and data reporting

Spearman’s rank-order correlation was used for all correlation statistics. A two-tailed Student’s *t*-test with *P* values adjusted for multiple testing (Bonferroni) was used to calculate the *P* values in **Fig. 1h-1i**, **Extended Data Fig. 6**, **Extended Data Fig. 7c**, and **Supplementary Table 3**. The error bars in all dot and bar plots show the standard deviation (s.d.) and were plotted with seaborn (0.10.0). The measure of center for the error bars is the mean. We did not predetermine sample sizes based on statistical methods. Investigators were not blinded to experimental conditions or assessment of experimental outcomes.

## Supporting information

Extended Data Figure 11 (part 1)

Extended Data Figure 11 (part 2)

## Acknowledgements

Support for this work was provided by the National Institutes of Health (R35 GM118158 and RM1 HG009490 to J.K.J.). J.K.J. was additionally supported by the Desmond and Ann Heathwood MGH Research Scholar Award and the Robert B. Colvin, M. D. Endowed Chair in Pathology. L.P. is supported by the National Human Genome Research Institute (NHGRI) Career Development Award (R00HG008399), Genomic Innovator Award (R35HG010717) and CEGS RM1HG009490. We thank Vikram Pattanayak and Julian Grunewald for discussions and technical advice and Ligi Paul Pottenplackel for assistance with editing the manuscript.

## Author contributions

J.Y.H., K.C.L., and J.Y.S. performed the laboratory experiments and J.Y.H. and J.Y.S. performed the computational analyses. J.Y.H., K.C.L., and J.K.J. conceived of and designed the study. J.K.J. and L.P. supervised the work. J.Y.H. and J.K.J. wrote the initial manuscript draft and all authors contributed to the writing of the final manuscript.

## Extended Data Figure Legends

**Extended Data Figure 1: Overview of prime editing**.

(**a**) The prime editor, a fusion protein of a CRISPR-SpCas9 H840A nickase and an engineered Moloney murine leukemia virus reverse transcriptase (MMLV-RT), coupled with a prime editing guide RNA (pegRNA) engages target DNA to form an R-loop and introduces a nick in the non-target strand (NTS). (**b**) The 3’ end of the NTS interacts with the 3’ end of the pegRNA extension, called the primer binding site (PBS), and this stabilized interaction allows for (**c**) extension of the NTS by reverse transcription with the reverse transcriptase and reverse transcription template (RTT) of the pegRNA where the edit of interest is encoded in the reverse transcribed RTT sequence. (**d**) The prime editor system dissociates from its target DNA and the newly synthesized edit-encoding “flap” strand competes with the wild-type (WT) strand for hybridization; flap excision and ligation occurs, with an optional target strand (TS) nick introduced with a nicking sgRNA (ngRNA) with the PE3 strategy. (**e**) The NTS sequence is preferentially repaired with the TS nick, resulting in a bias towards successful editing in scenarios where the edit-encoding flap outcompetes the WT flap for genomic incorporation. Figure adapted from Hsu et al. 2021^15^.

**Extended Data Figure 2: Saturation mutagenesis efficiencies for single-base and triple-base substitutions**.

Single-base (**a,c,e**) and triple-base (**b,d,f**) saturation mutagenesis efficiencies across RNF2 site 1, RUNX1 site 1, and VEGFA site 4. Data shown are from three independent biological replicates, meanslJ±lJs.d. are indicated. Sequences of oligonucleotides used to construct the pegRNAs are provided in **Supplementary Table 1**.

**Extended Data Figure 3: Installation of randomized insertion edits at genomic targets**.

Editing efficiencies (**a**) and the total count of unique insertion edits across replicates (**b**) for randomized insertion edits of various lengths (3, 6, and 9 bp) at HEK site 3, RNF2 site 1, RUNX1 site 1, and VEGFA site 4. Data shown are from three independent biological replicates, meanslJ±lJs.d. are indicated. Sequences of oligonucleotides used to construct the pegRNAs are provided in **Supplementary Table 1**.

**Extended Data Figure 4: Screening of pegRNA-ngRNA combinations at the *BCR-ABL1* locus in HEK293T cells**.

Screening of 84 pegRNA-ngRNA combinations to install a randomized (NNN) substitution edit at amino acid position 315 in *BCR-ABL1*. Experiments were performed in HEK293T cells. Data shown are from three independent biological replicates, meanslJ±lJs.d. are indicated. Sequences of oligonucleotides used to construct the pegRNAs are provided in **Supplementary Table 2**.

**Extended Data Figure 5: Overview of saturation mutagenesis efficiencies at BCR-ABL1 with and without imatinib drug treatment in K562 cells**.

(**a**) Saturation mutagenesis efficiencies at BCR-ABL1 in K562 cells following 7 days of imatinib or DMSO treatment. Correlation (Spearman’s Rho) between (**b**) DMSO and (**c**) imatinib treated samples for the count of variant NNK codons across amino acid positions 311-318 at BCR-ABL1. Saturation mutagenesis efficiencies by amino acid position for (**d,f**) DMSO and (**e,g**) imatinib treated samples.

**Extended Data Figure 6: Count of NNK codons across saturation mutagenesis window at *BCR-ABL1* locus**.

Count of installed NNK codons across amino acid positions 311-318 within *BCR-ABL1* for (**a,b**) DMSO and (**c,d**) imatinib treated samples. Certain NNK codons at specific amino acid positions are labeled as WT because they are the reference sequence and therefore aren’t counted as installed edits.

**Extended Data Figure 7: Enrichment scores for amino acid variants from the *BCR-ABL1* imatinib resistance screen in K562 cells.**

Enrichment scores for all amino acid variants within residue positions 311-318 in *BCR-ABL1* associated with imatinib resistance in K562 cells. Statistical significance was determined by a t-test comparing any given amino acid variant against a null distribution consisting of NNK codons coding for silent mutations within the BCR-ABL1 protein, followed by multiple testing correction using the Bonferroni method. Statistics are provided in **Supplementary Table 3**.

**Extended Data Figure 8: Application of MOSAIC for the fine-mapping of non-coding regulatory sequences within the *IRF1* 5’ UTR**.

(**a**) Schematic for *in situ* saturation mutagenesis of the IRF1 5’ UTR to discover non-coding regulatory sequence elements involved in its transcription regulation in HEK293T cells. (**b**) Screening of 112 pegRNA-ngRNA combinations to install randomized substitution edits across the *IRF1* 5’ UTR. (**c**) Enrichment scores of 3 bp tiles across the IRF1 5’ UTR sequence associated with *IRF1* transcription (mRNA vs. gDNA). Statistical significance was determined by a t-test comparing any given 3-bp mutagenized tile against a null distribution consisting of the reference *IRF1* 5’UTR sequence, followed by multiple testing correction using the Bonferroni method. Data shown are from three independent biological replicates, meanslJ±lJs.d. are indicated. Sequences of oligonucleotides used to construct the pegRNAs are provided in **Supplementary Table 4**.

**Extended Data Figure 9: Correlation between pooled pegRNA optimizations and individual pegRNA optimizations performed in Anzalone et al. 2019**.

Correlation (Spearman’s Rho) between pooled pegRNA optimizations with MOSAIC and individual pegRNA optimizations performed in Anzalone et al. 2019 for EMX1 site 1, HEK site 3, and RNF2 site 1 genomic targets in HEK293T cells.

**Extended Data Figure 10: Mean prime editing frequency with various PBS and RTT lengths from pegRNA design optimization**.

The mean prime editing frequency using pegRNAs with various length (**a**) PBS (8-17 nts) and (**b**) RTT (10-30 nts) sequences from pegRNA optimization profiles 89 target spacer sequences.

**Extended Data Figure 11: pegRNA optimization profiles across 89 spacer sequences**.

Averaged PE efficiencies for 89 target spacer sequences spanning all 210 combinations of PBS (8-17 nts) and RTT (10-30 nt) lengths used the pegRNA pools.

## Supplementary Notes

**Supplementary Note 1: PCR-generated prime editing guide RNAs (pegRNAs)**

**Supplementary Note 2: High-throughput pegRNA optimization with MOSAIC**

## Supplementary Table legends

**Supplementary Table 1:** Oligonucleotide sequences to construct PCR pegRNAs and ngRNAs for saturation mutagenesis and randomized insertions at HEK site 3, RNF2 site 1, RUNX1 site 1, and VEGFA site 4

**Supplementary Table 2:** Oligonucleotide sequences to construct PCR pegRNAs and ngRNAs for saturation mutagenesis at the BCR-ABL1 imatinib binding site

**Supplementary Table 3:** Statistics for amino acid variation in imatinib resistance *BCR-ABL1* mutagenesis screen

**Supplementary Table 4:** Oligonucleotide sequences to construct PCR pegRNAs and ngRNAs for saturation mutagenesis at the IRF1 5’ UTR element

**Supplementary Table 5:** Oligonucleotide sequences to construct PCR pegRNAs and ngRNAs to assess barcoding effects for pooled pegRNA optimization

**Supplementary Table 6:** Oligonucleotide sequences to construct PCR pegRNAs and ngRNAs to assess efficiencies of pooled and single construct prime editing

**Supplementary Table 7:** Target sites for high-throughput pooled pegRNA optimization

**Supplementary Table 8:** Oligonucleotide sequences to perform large-scale high-throughput pooled pegRNA optimization

**Supplementary Table 9:** Highest activity, highest averaged activity, and 25th percentile pegRNA designs tested

**Supplementary Table 10:** NGS primer sequences for pegRNA optimizations

## References

1. Anzalone, A. V. et al. Search-and-replace genome editing without double-strand breaks or donor DNA. Nature 576, 149–157 (2019).

2. Cong, L. et al. Multiplex genome engineering using CRISPR/Cas systems. Science 339, 819–823 (2013).

3. Mali, P. et al. RNA-guided human genome engineering via Cas9. Science 339, 823–826 (2013).

4. Hwang, W. et al. Efficient genome editing in zebrafish using a CRISPR-Cas system. Nat. Biotechnol. 31, 227–229 (2013).

5. Komor, A. C. et al. Programmable editing of a target base in genomic DNA without double-stranded DNA cleavage. Nature 533, 420–424 (2016).

6. Gaudelli, N. M. et al. Programmable base editing of AT to GC in genomic DNA without DNA cleavage. Nature 551, 464–471 (2017).

7. Kurt, I. C., et al. CRISPR C-to-G base editors for inducing targeted DNA transversions in human cells. Nat. Biotechnol. 39, 41–46 (2021).

8. Chen, et al. Enhanced prime editing systems by manipulating cellular determinants of editing outcomes. Cell 184 (22), 5635–5652 (2021).

9. Nelson, J. W. et al. Engineered pegRNAs improve prime editing efficiency. Nat. Biotechnol. 40, 402–410 (2022).

10. Ren, R. et al. Mechanisms of BCR–ABL in the pathogenesis of chronic myelogenous leukaemia. Nature Reviews Cancer 5, 172–183 (2005).

11. Weisberg, E. et al. Resistance to imatinib (Glivec): update on clinical mechanisms. Drug Resistance Updates 6, 231–238 (2003).

12. Drullion, C. et al. Apoptosis and autophagy have opposite roles on imatinib-induced K562 leukemia cell senescence. Cell Death & Disease 3, 373 (2012).

13. Ma, L. et al. CRISPR-Cas9–mediated saturated mutagenesis screen predicts clinical drug resistance with improved accuracy. PNAS 114 (44), 11751–11756 (2017).

14. Burke, T. W. et al. The downstream core promoter element, DPE, is conserved from Drosophila to humans and is recognized by TAFII60 of Drosophila. Genes & Development 11, 3020–3031 (1997).

15. Hsu, J. Y. et al. PrimeDesign software for rapid and simplified design of prime editing guide RNAs. Nat. Comm. 12, 1034 (2021).

16. Kim, H. K. et al. Predicting the efficiency of prime editing guide RNAs in human cells. Nat. Biotechnol. 39, 198–206 (2021).

17. Erwood, S. et al. Saturation variant interpretation using CRISPR prime editing. Nature Biotechnology, 40 (6), 885–895 (2022).

18. Rees, H. et al. Base editing: precision chemistry on the genome and transcriptome of living cells. Nature Reviews Genetics, 19 (12), 770–788 (2018).

19. Anzalone, A. V. et al. Programmable deletion, replacement, integration and inversion of large DNA sequences with twin prime editing. Nat. Biotechnol. 40, 731–740 (2022).

