## Supplementary figures and images for "MOSAIC enables *in situ* saturation mutagenesis of genes and CRISPR prime editing guide RNA optimization in human cells"

### Extended Data Figure 1

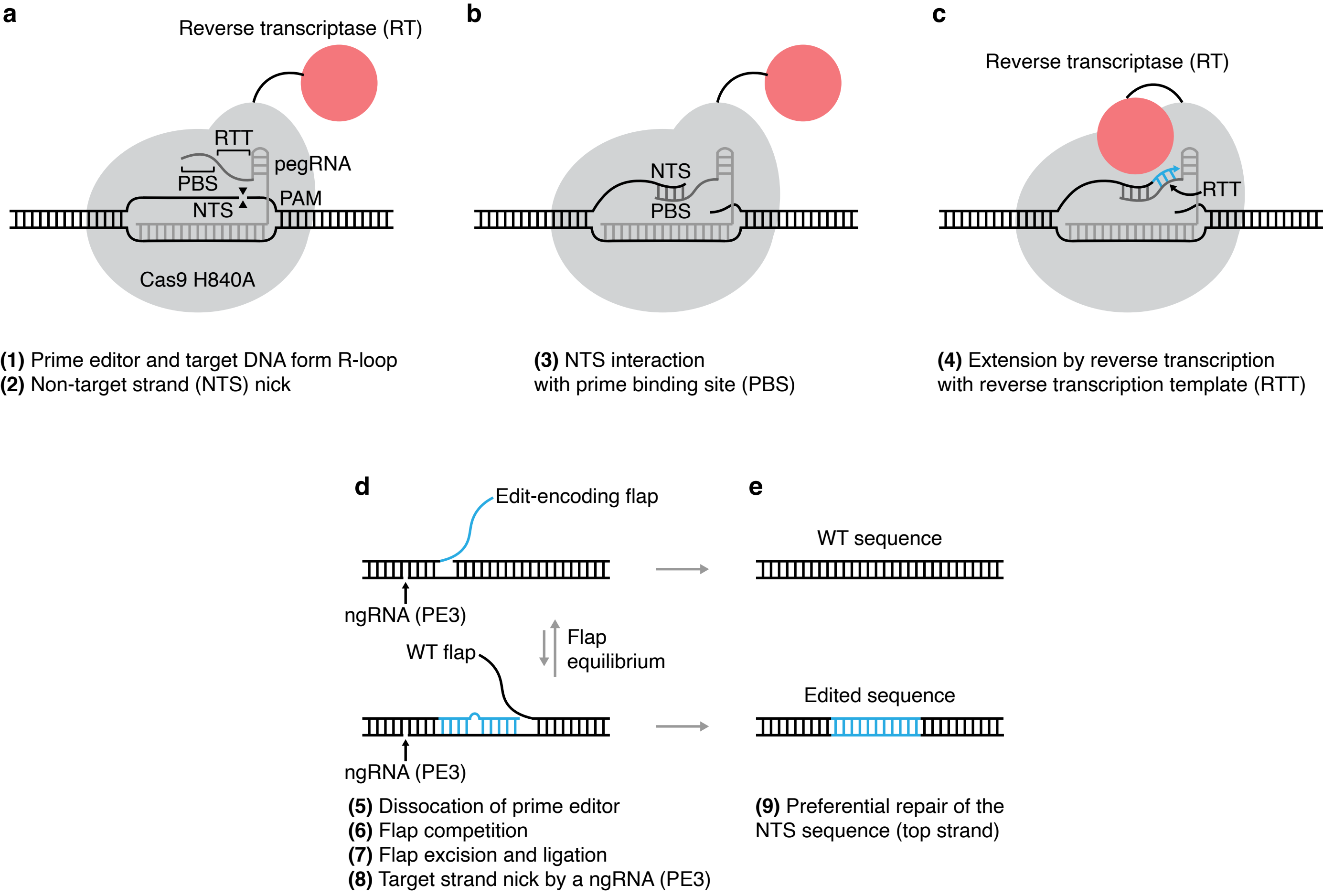

### Extended Data Figure 2

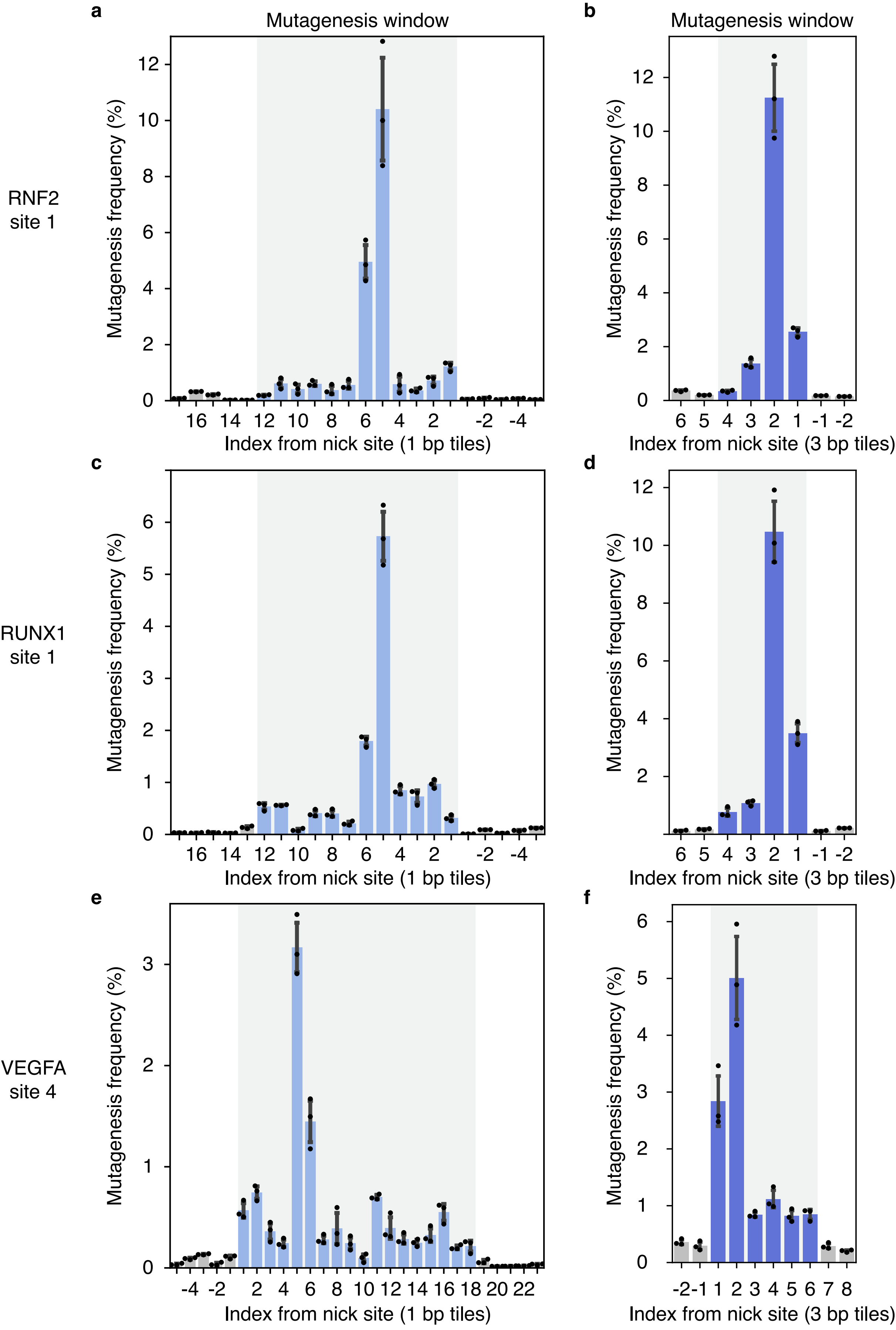

### Extended Data Figure 3

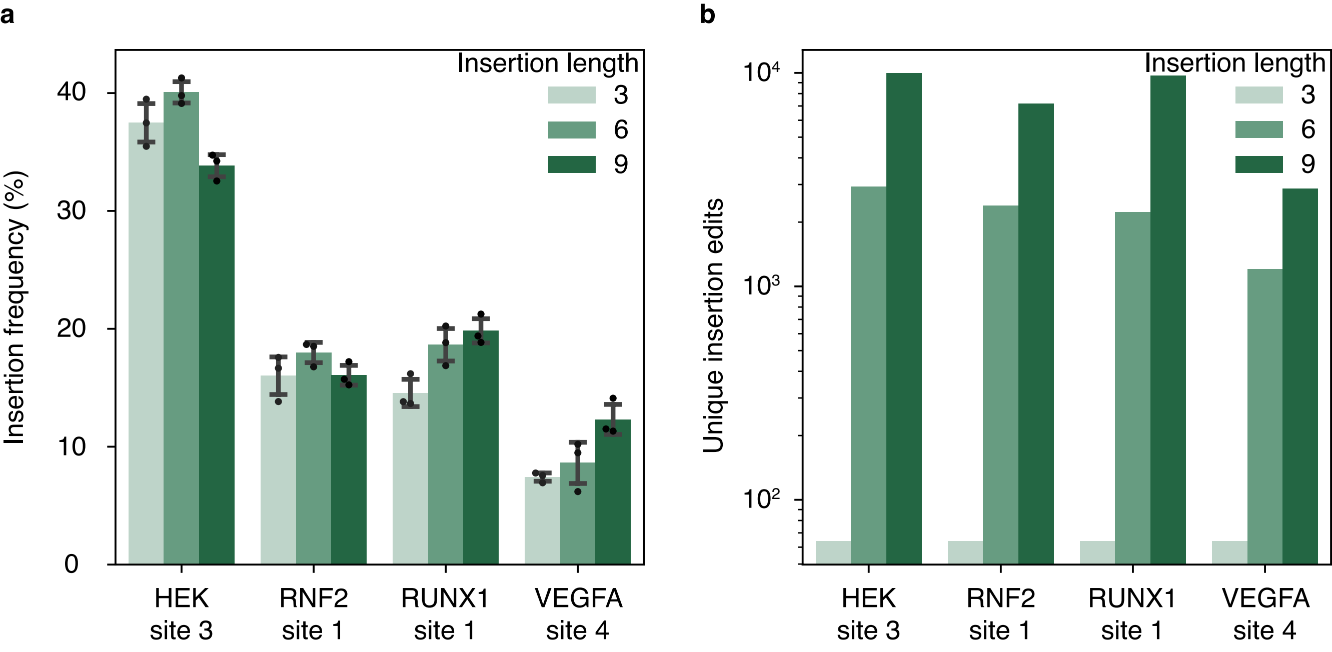

### Extended Data Figure 4

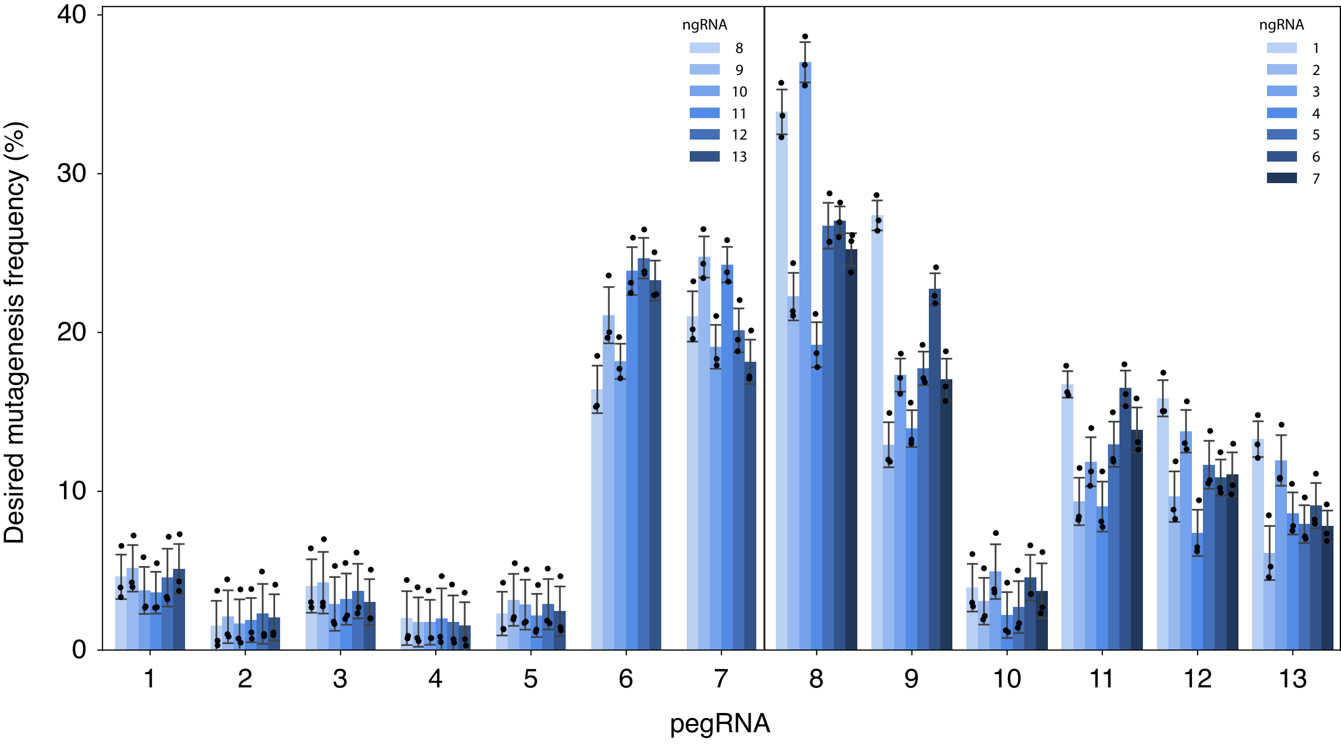

### Extended Data Figure 5

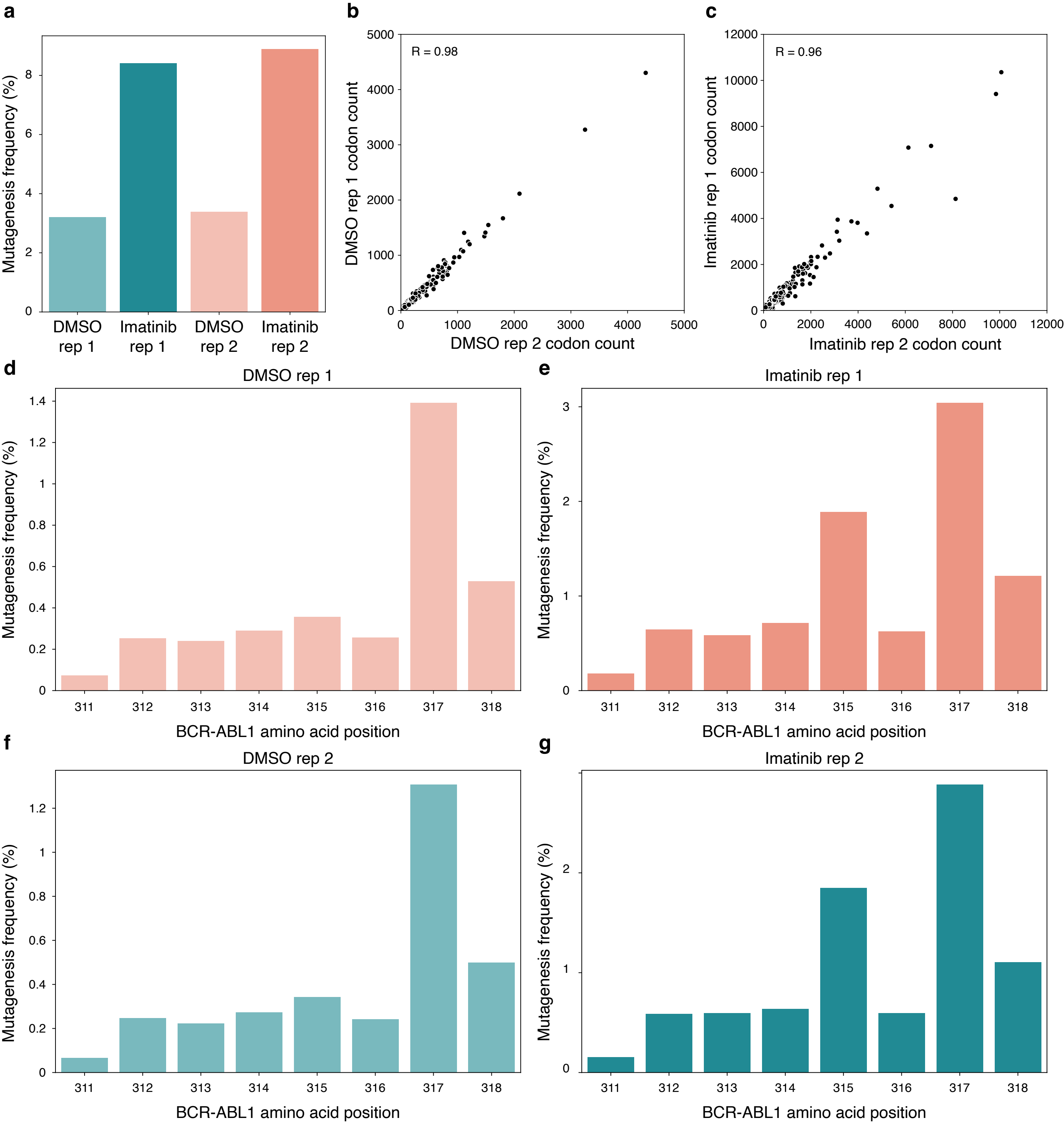

### Extended Data Figure 6

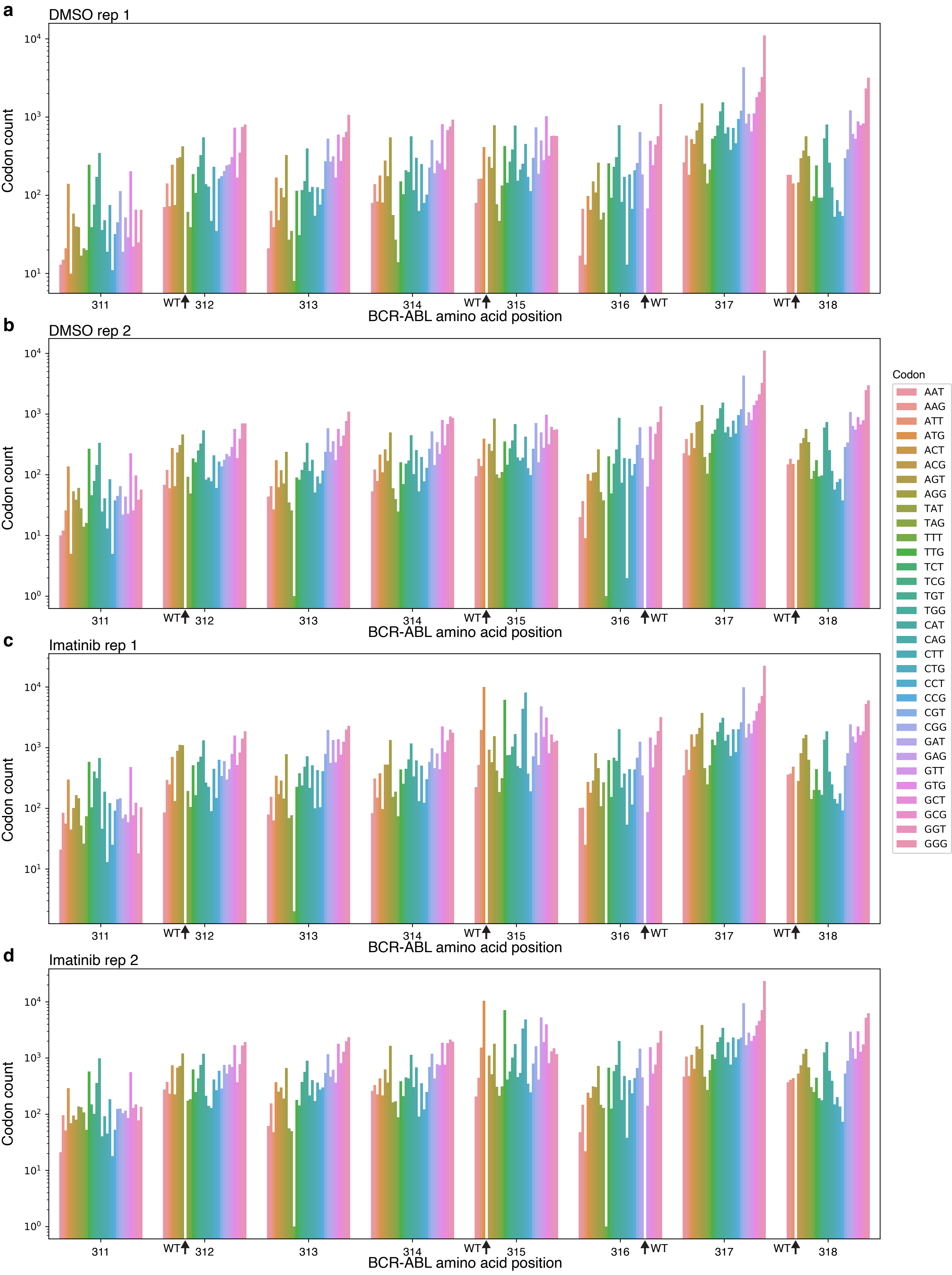

### Extended Data Figure 7

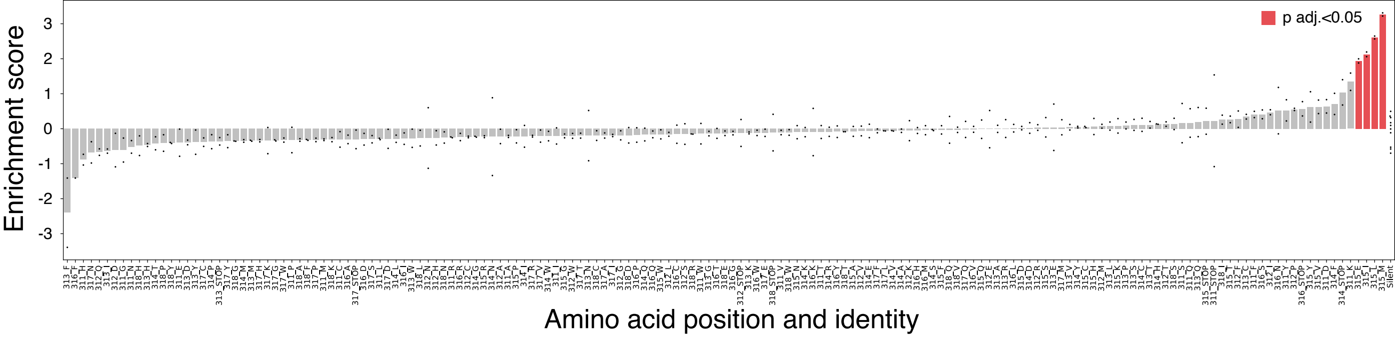

### Extended Data Figure 8

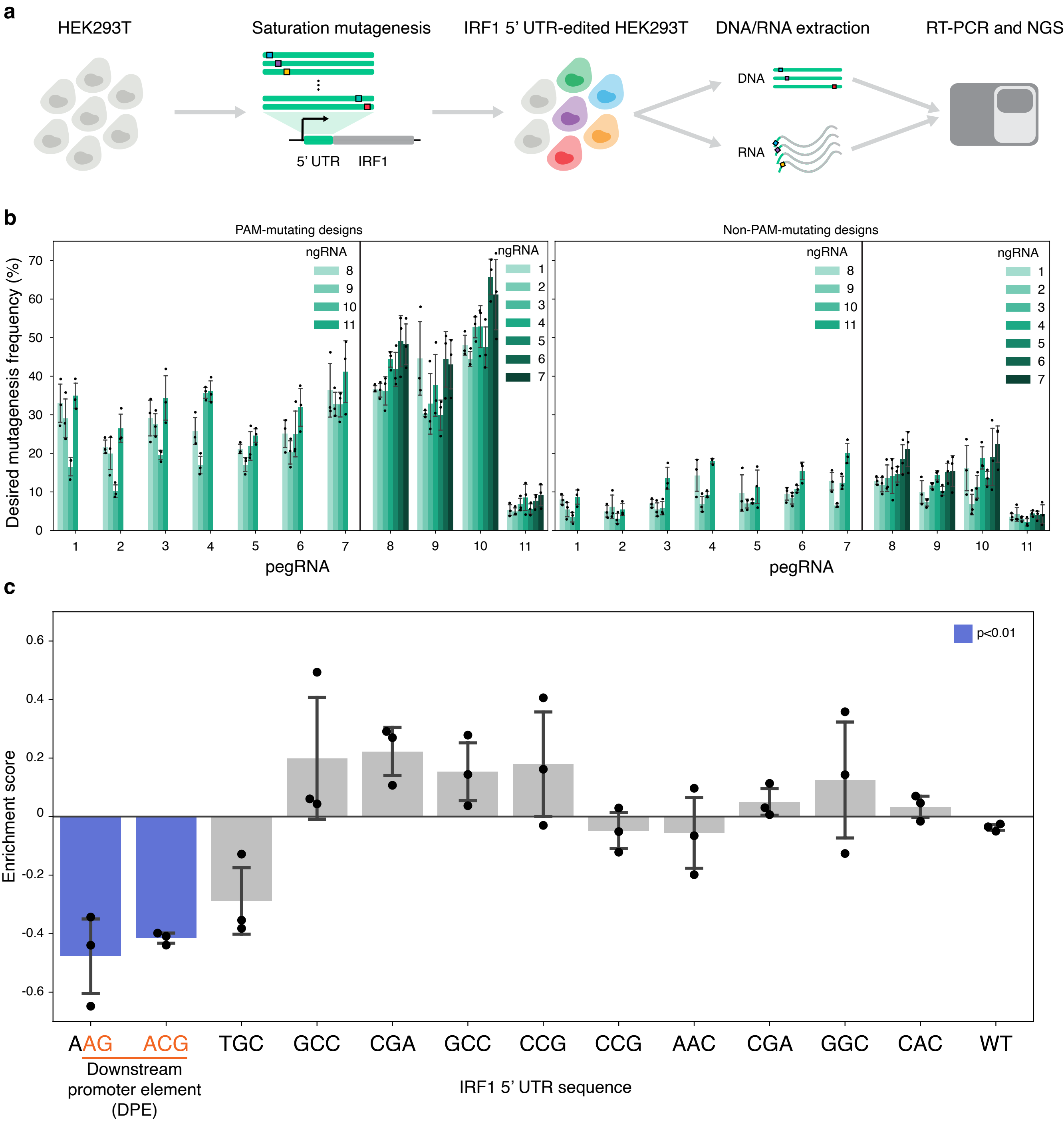

### Extended Data Figure 9

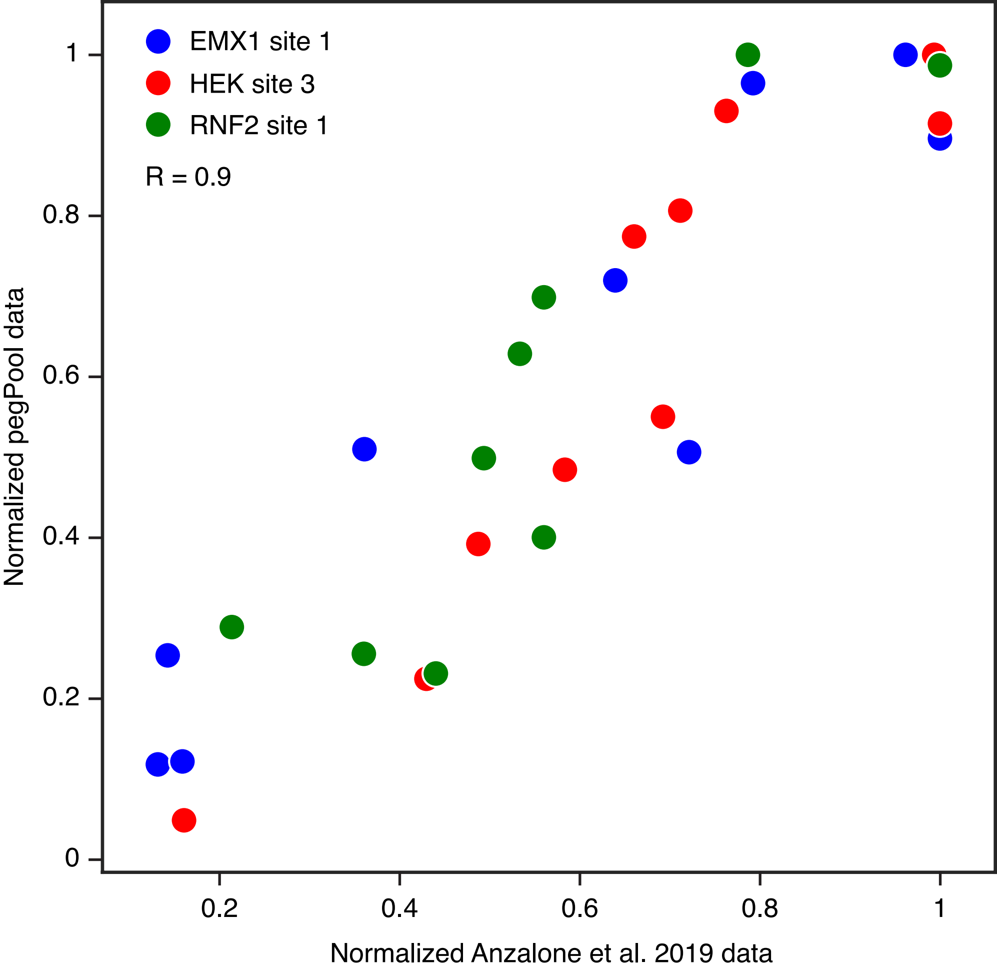

### Extended Data Figure 10

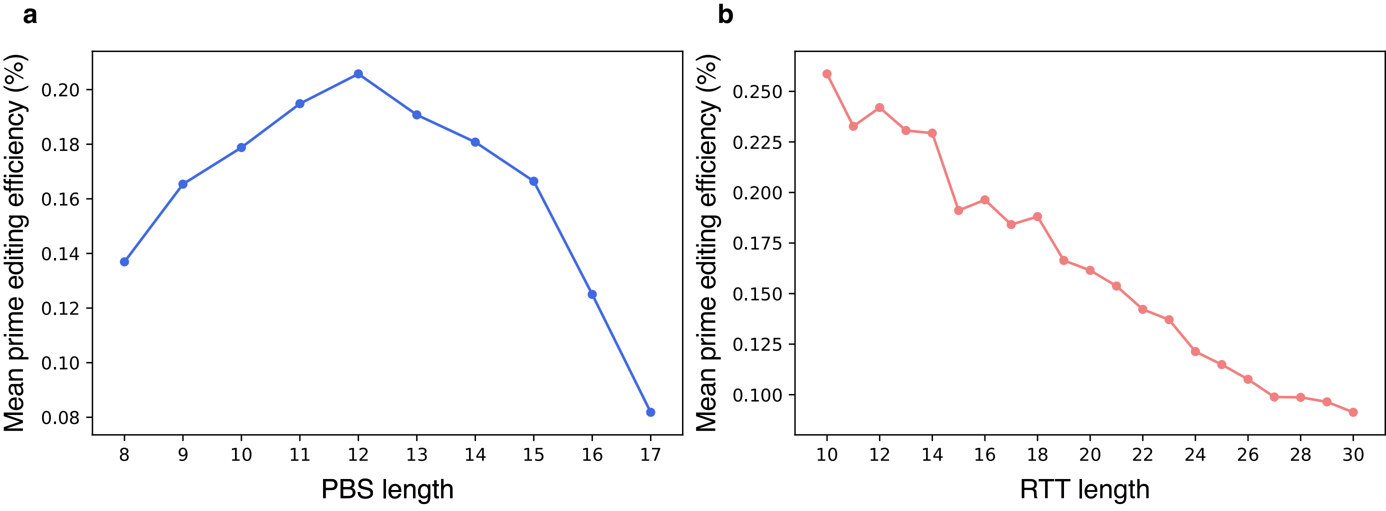

### Extended Data Figure 11 (part 1)

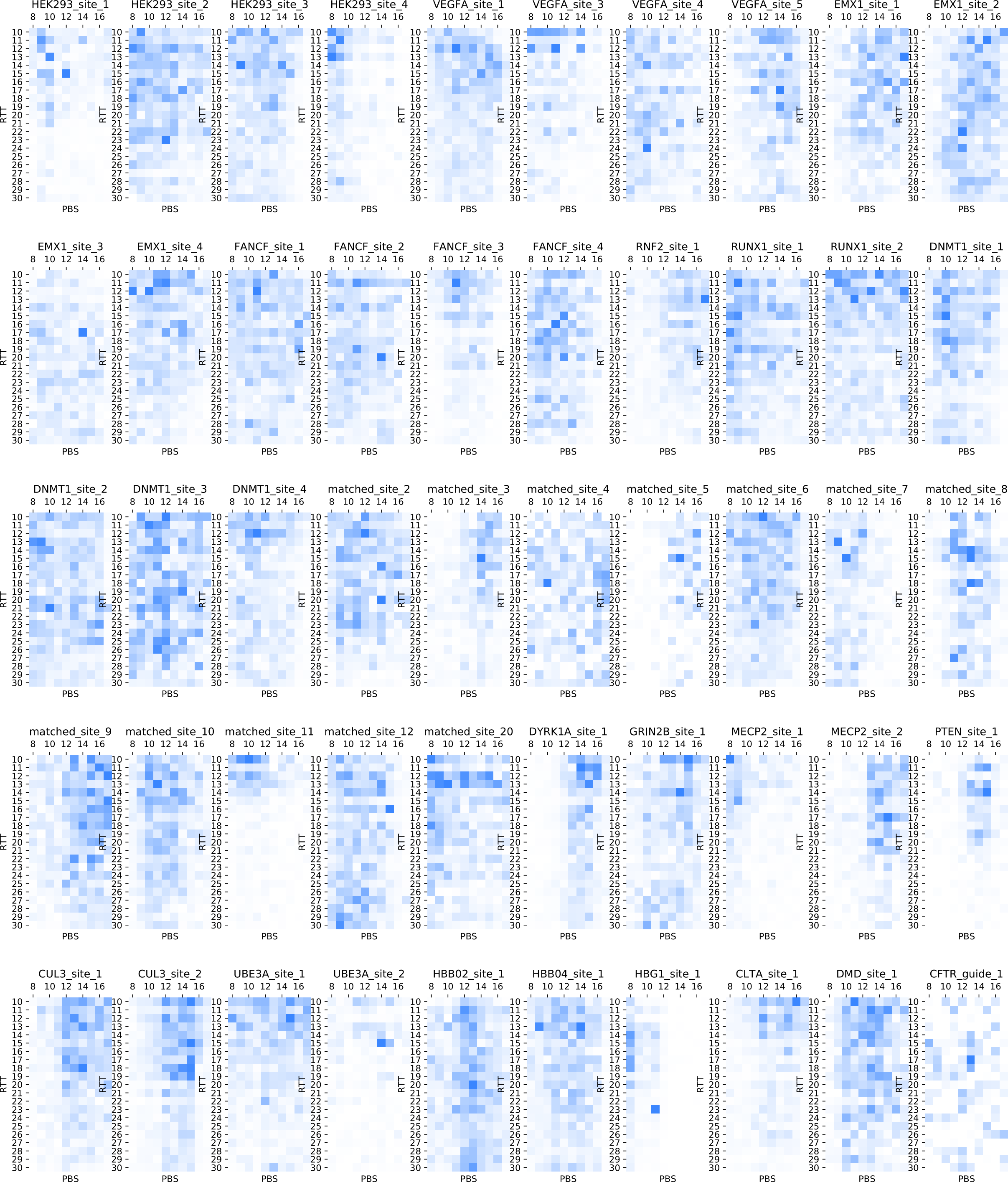

### Extended Data Figure 11 (part 2)

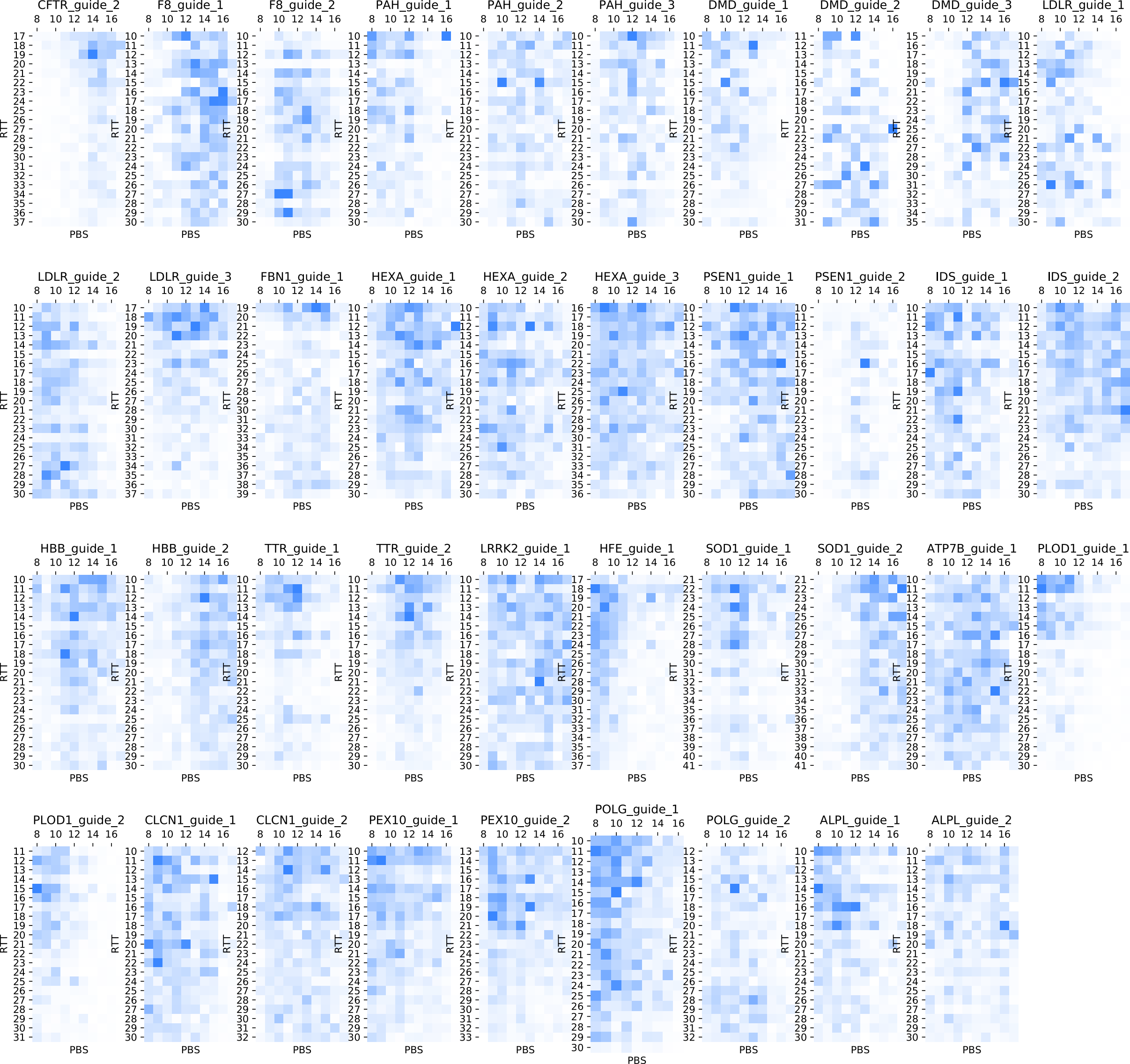
