## Supplementary Notes for "MOSAIC enables *in situ* saturation mutagenesis of genes and CRISPR prime editing guide RNA optimization in human cells"

**Supplementary Note 1: PCR-generated prime editing guide RNAs (pegRNAs)**

The construction of pegRNAs or ngRNAs by PCR requires two sequential steps. The first PCR steps (labeled PCR1 below) are for the amplification of the “U6 promoter PCR fragment”, “pegRNA PCR fragment”, and “ngRNA PCR fragment”. Following the completion of these first PCR reactions, the desired amplification products are cleaned and isolated with paramagnetic beads and two 75% ethanol washes. The second PCR steps (labeled PCR2 below) fuse the “U6 promoter PCR fragment” to either the “pegRNA PCR fragment” or “ngRNA PCR fragment” to obtain pegRNA and ngRNA constructs, respectively. Following the completion of these second PCR reactions, the desired amplification products are cleaned and isolated with paramagnetic beads and two 75% ethanol washes. These purified PCR products are ready for transfection into mammalian cells. Standard PCR protocol is recommended based on the PCR kit used.

**-----------------------------------------------------------PCR1-----------------------------------------------------------**

PCR1 reaction to amplify “U6 promoter PCR fragment”

| **Component** | **Volume (µL)** |
| --- | --- |
| 5X PCR Buffer | 10 |
| 10 mM dNTPs | 1 |
| 10 µM *U6 forward primer* | 2.5 |
| 10 µM *U6 reverse primer* | 2.5 |
| U6 promoter-containing plasmid (1-5 ng/uL) | 1 |
| Polymerase | 0.5 |
| Water | 32.5 |
| Total | 50 |

*Clean up with 0.7X paramagnetic beads

PCR1 reaction to assemble “pegRNA PCR fragment”

| **Component** | **Volume (µL)** |
| --- | --- |
| 5X PCR Buffer | 10 |
| 10 mM dNTPs | 1 |
| 10 µM *pegRNA forward primer* | 2.5 |
| 10 µM *pegRNA reverse primer* | 2.5 |
| 10 µM *tracrRNA oligo* | 1 |
| 10 µM *Spacer oligo* | 1 |
| 10 µM *pegRNA extension oligo 1* | 1 |
| 10 µM *pegRNA extension oligo 2 (optional)* | 0 |
| Polymerase | 0.5 |
| Water | 30.5 |
| Total | 50 |

*Clean up with 1.2X paramagnetic beads

PCR1 reaction to assemble “ngRNA PCR fragment”

| **Component** | **Volume (µL)** |
| --- | --- |
| 5X PCR Buffer | 10 |
| 10 mM dNTPs | 1 |
| 10 µM *ngRNA forward primer* | 2.5 |
| 10 µM *ngRNA reverse primer* | 2.5 |
| 10 µM *tracrRNA oligo* | 1 |
| 10 µM *Spacer oligo* | 1 |
| Polymerase | 0.5 |
| Water | 31.5 |
| Total | 50 |

*Clean up with 1.2X paramagnetic beads

**-----------------------------------------------------------PCR2-----------------------------------------------------------**

PCR2 reaction to fuse “U6 promoter PCR fragment” with “pegRNA PCR fragment”

| **Component** | **Volume (µL)** |
| --- | --- |
| 5X PCR Buffer | 10 |
| 10 mM dNTPs | 1 |
| 10 µM *U6 forward primer* | 2.5 |
| 10 µM *pegRNA reverse primer* | 2.5 |
| “U6 promoter PCR fragment” (1-5 ng/uL) | 1 |
| “pegRNA PCR fragment” (1-5 ng/uL) | 1 |
| Polymerase | 0.5 |
| Water | 31.5 |
| Total | 50 |

*Clean up with 0.7X paramagnetic beads

PCR2 reaction to fuse “U6 promoter PCR fragment” with “ngRNA PCR fragment”

| **Component** | **Volume (µL)** |
| --- | --- |
| 5X PCR Buffer | 10 |
| 10 mM dNTPs | 1 |
| 10 µM *U6 forward primer* | 2.5 |
| 10 µM *ngRNA reverse primer* | 2.5 |
| “U6 promoter PCR fragment” (1-5 ng/uL) | 1 |
| “pegRNA PCR fragment” (1-5 ng/uL) | 1 |
| Polymerase | 0.5 |
| Water | 31.5 |
| Total | 50 |

*Clean up with 0.7X paramagnetic beads

**Oligonucleotide and primer sequences**

*U6 forward primer: CTGTACAAAAAAGCAGGCTTTAAAGGAACCAATTC*

*U6 reverse primer: GGTGTTTCGTCCTTTCCACAAGATATATAAAGC*

*pegRNA forward primer: GCTTTATATATCTTGTGGAAAGGACGAAACACC*

*pegRNA reverse primer: GCAGCACGTGATACACCAAAAAAA*

*ngRNA forward primer: GCTTTATATATCTTGTGGAAAGGACGAAACACC*

*ngRNA reverse primer: GCAGCACGTGATACACCAAAAAAAGCACCGACTCGGTGCC*

*Spacer oligo*

*ATATCTTGTGGAAAGGACGAAACACCNNNNNNNNNNNNNNNNNNNNGTTTTAGAGCTAGAAATAGCAAGTTAAAATAAG*

where the Nx20 sequence represents the spacer sequence. Make sure the 5’ element is a G base for efficient transcription off the U6 promoter.

*tracrRNA oligo*

*GCACCGACTCGGTGCCACTTTTTCAAGTTGATAACGGACTAGCCTTATTTTAACTTGCTATTTCTAGCTCTAAAAC*

*pegRNA extension oligo 1*

*AAGTGGCACCGAGTCGGTGC + [insert pegRNA 3’ extension sequence 5’ to 3’] + TTTTTTTGGTGTATCACGTGCTGC*

Depending on the length of the pegRNA extension, additional pegRNA extension oligos may be needed to assemble the full pegRNA 3’ extension. For example, if the pegRNA extension is too long to fit on a single oligo and requires two pegRNA extension oligos, it would take on the following form:

*pegRNA extension oligo 1*

*AAGTGGCACCGAGTCGGTGC + [insert pegRNA 3’ extension sequence 5’ to 3’]*

*pegRNA extension oligo 2*

*GCAGCACGTGATACACCAAAAAAA + [insert reverse complement pegRNA 3’ extension sequence 5’ to 3’]*

**Important note**: The *pegRNA extension oligos 1 and 2* will require some degree of sequence overlap at the 3’ end to mediate successful PCR assembly. We recommend at least 15 bp of overlap (or sufficient melting temperature of ~60°C).

**Supplementary Note 2: High-throughput pegRNA optimization with MOSAIC**

We designed and synthesized an oligonucleotide library containing 21,210 members (Supplementary Table 8) to perform pegRNA optimizations across 89 different spacer sequences. Each library member contained a spacer sequence, tracrRNA sequence, and pegRNA 3’ extension of varied length. For each spacer sequence, a total of 210 PBS-RTT length combinations were designed for the 3’ extension with PBS lengths spanning 8-17 nt and RTT lengths typically spanning 10-30 nt, where each of these PBS-RTT length combinations were uniquely barcoded with a 4-mer insertion edit placed at the +1 position. Unique flanking sequences were used to amplify sub-libraries from the oligonucleotide pool corresponding to a specific pool of pegRNA designs for a single spacer sequence. Following low-cycle PCR amplification of the sub-libraries, PCR assembly was used to fuse a U6 promoter element to each of the sub-libraries for expression in human cells (Supplementary Note 1). These pegRNA optimization sub-libraries were then transfected into human cells with PE2 plasmid to perform pooled pegRNA optimizations.

Following transfection of the pegRNA optimization pegPools into HEK293T cells and subsequent NGS analysis, we identified top-performing pegRNA designs for each spacer sequence based on the frequency of the 4-mer insertion barcode edits. The ‘highest activity’ pegRNA design refers to the individual PBS-RTT length combination barcode that was observed with highest frequency in the next-generation sequencing data. The ‘highest averaged activity’ pegRNA design utilizes averaged information from an individual pegRNA optimization profile. The ‘highest averaged activity’ PBS length is chosen by averaging all values across different RTT lengths for a given PBS length for all the different PBS lengths, and then choosing the PBS length with the highest value. The ‘highest averaged activity’ RTT length is chosen by averaging all values across different PBS lengths for a given RTT length for all the different RTT lengths, and then choosing the RTT length with the highest value. The 25^th^ percentile pegRNA designs refer to designs ranked at the 25^th^ percentile within the 210 different pegRNA designs for a given spacer sequence. To validate these identified pegRNA designs, we installed all possible single base substitutions and a variety of small insertion and deletion mutations across 20 different spacer sequences.
